## Supplemental Figures for "Nerve Growth Factor Receptor Limits Inflammation to Promote Remodeling and Repair of Osteoarthritic Joints"

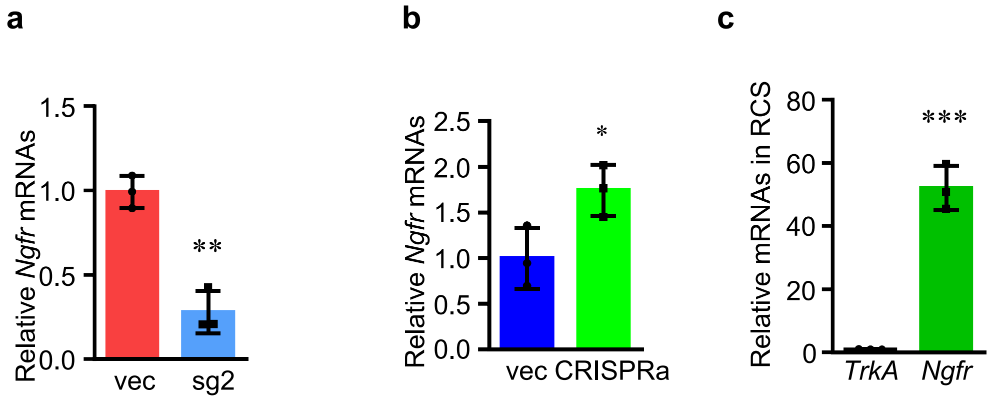


**Supplementary Figure 1.** Quantitative RT-PCR of *Ngfr* and *TrkA* mRNAs. **a**, CRISPR-mediated deletion of *Ngfr* in C2C12 cells. vec, empty vector; sg2, single guide RNAs #2 targeting *Ngfr*. **b**, CRISPR-mediated transcriptional activation of *Ngfr*. **c**, Comparison of *TrkA* and *Ngfr* in RCS cells. n = 3. *p<0.05, **p<0.01, ***p<0.001.

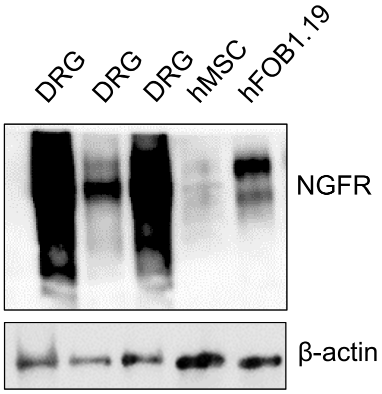


**Supplementary Figure 2.** The protein levels in dorsal root ganglions, mesenchymal stem cells, and osteoblasts, as shown by western blot of NGFR proteins extracted from the indicated tissues or cells.

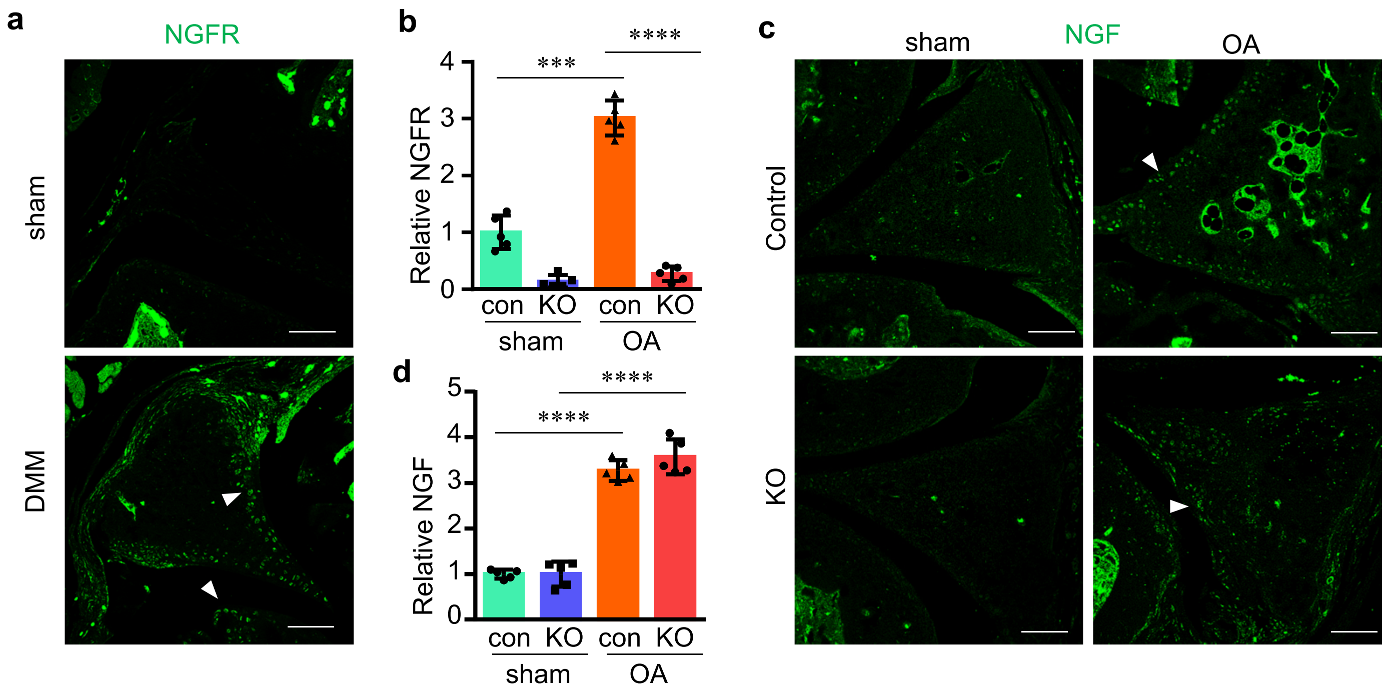


**Supplementary Figure 3.** NGFR and NGF are induced in osteoarthritic joints. **a**, NGFR induction in the knee joints 4 weeks after the DMM surgery. Arrowheads, NGFR-positive IHC signals. **b**, Quantification of NGFR IHC results shown in Fig. 3a. Control, *Ngfr* floxed. KO, $Ngfr^{Agc1\text{-}CreER}$ mice injected with tamoxifen weekly to ablate *Ngfr*, starting from 10 days after the DMM surgery. **c** and **d**, NGF is induced in osteoarthritic joints. Arrowheads: $NGF^{+}$ osteochondral cells. Scale bar, 100 μm. n = 5. ***p<0.001, ****p<0.0001.

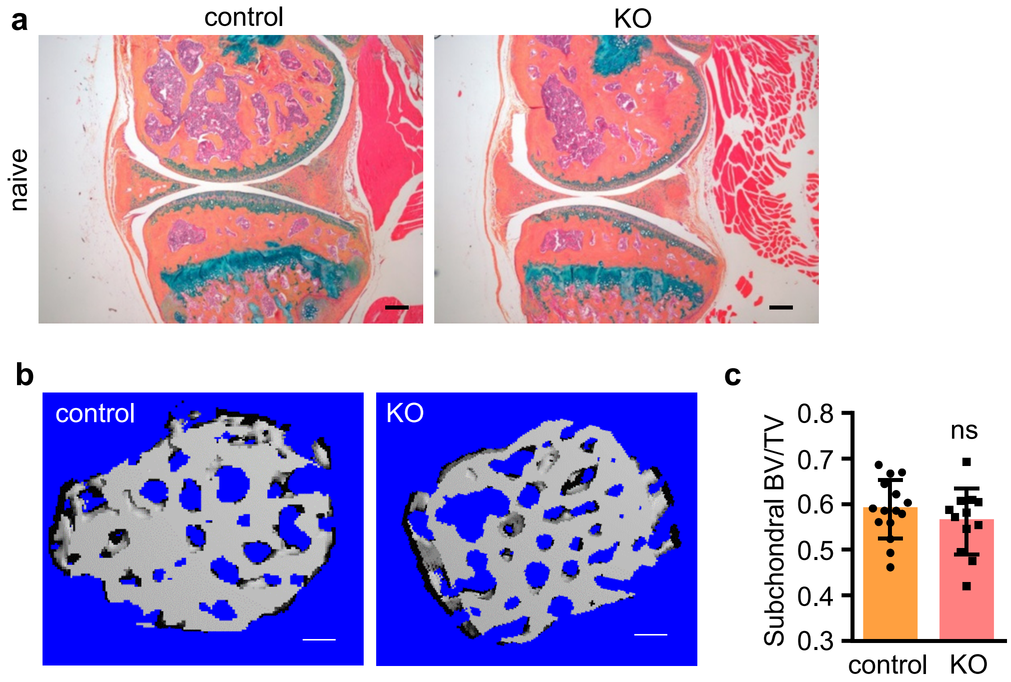


**Supplementary Figure 4.** NGFR ablation in osteochondral cells induced in pre-adulthood may not dysregulate joint development and homeostasis. **a**, Representative histology images of knee joint sections stained by Alcian blue/hematoxylin & orange G. The joints were collected 5 months after 5 consecutive days of injection, which was performed on the mice aged 15 days old. Scale bar: 200 μm. **b** and **c**, The subchondral bone mass in the non-osteoarthritic knees of control and NGFR KO mice (6 months old) does not show significant differences. **b**, Representative μCT 3D images of subchondral bone. **c**, Quantification of subchondral bone mass (BV/TV). n ≥ 12. Scale bar: 150 μm.

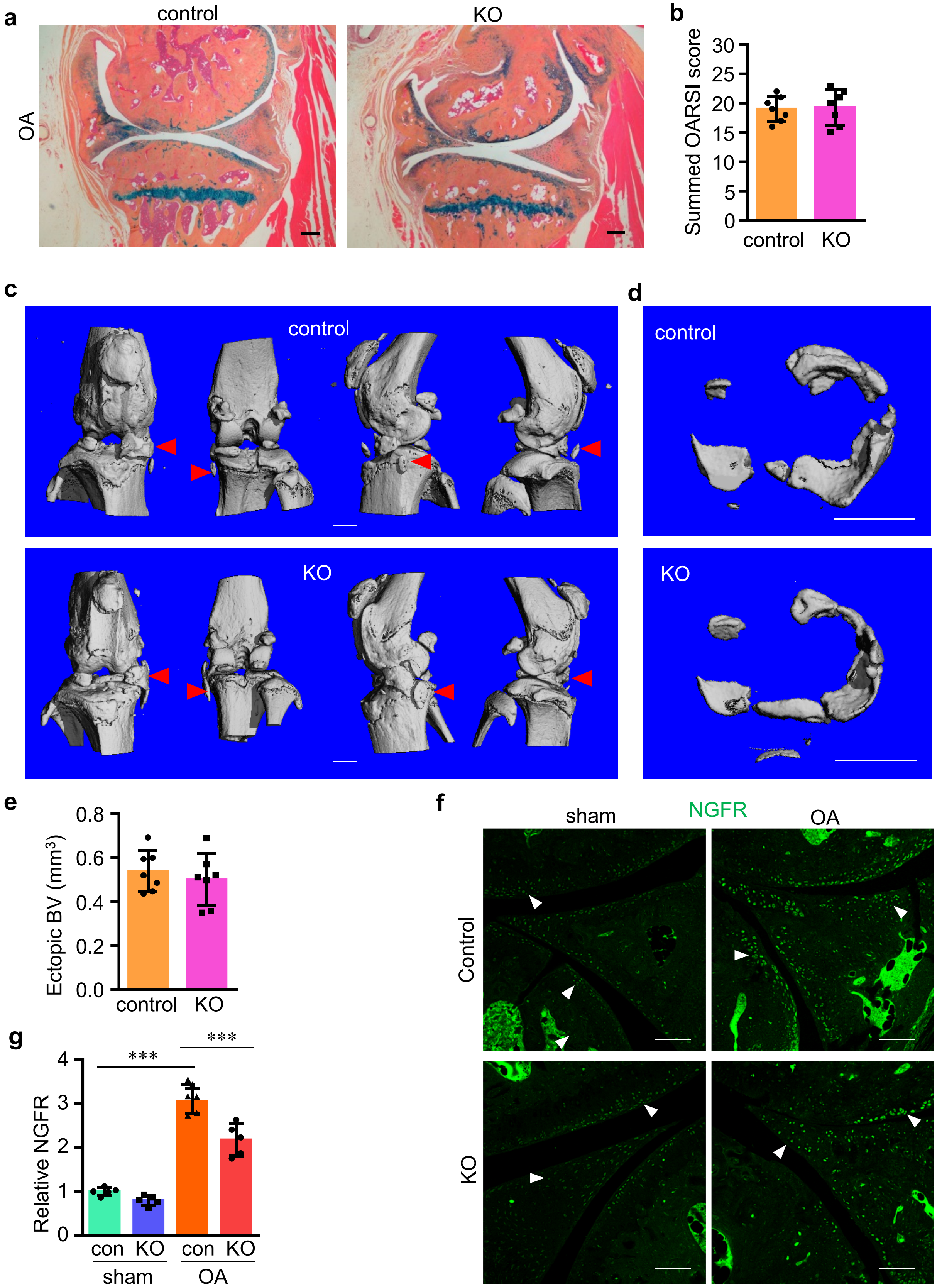


**Supplementary Figure 5.** Pre-adulthood loss-of-function of NGFR in osteochondral cells has no significant effects on OA progression. The mice were injected with tamoxifen for 5 consecutive days at the age of 15 days old. When they were 3 months old, the DMM surgery was performed on their right knees to induce OA. **a**, Representative histology staining with alcian blue/hematoxylin & orange G. Scale bar: 200 µm. **b**, OARSI scoring of histology results of OA joints. **c**, Representative µCT 3D images of control and KO joints with OA . Red arrowheads, ectopic bone outgrowth. **d** and **e**, Representative µCT 3D images and quantification of ectopic bone growth around meniscus and synovium as well as quantification of ectopic BV. Scale bars: 1 mm. n=7. **f** and **g**, NGFR ablation is not substantial in the joints of KO mice receiving tamoxifen at young ages. Arrowheads: NGFR+ osteochondral cells. Scale bar, 100 µm. n = 5. ***p<0.001.

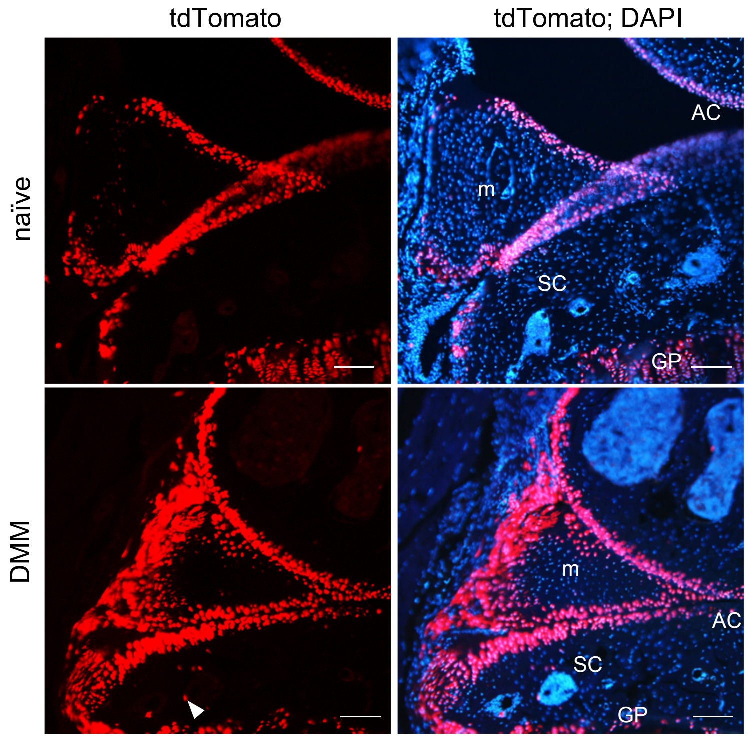


**Supplementary Figure 6.** *Aggrecan-CreER* efficiently targets multiple osteochondral tissues in the osteoarthritic (DMM) joints, including articular cartilage (AC), subchondral area (SC), meniscus (m), and targets mainly cartilaginous tissues in unoperated (naïve) knee joints. Arrowhead, targeted cells in SC. Growth plate (GP). n=3. Scale bar, 100 µm.

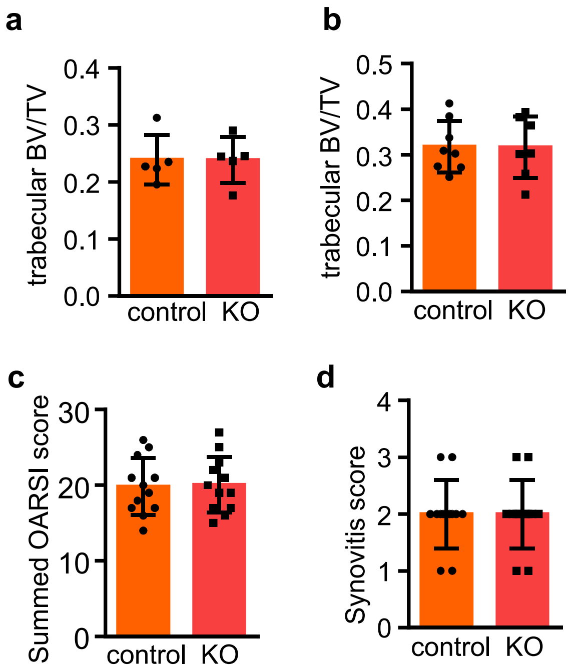


**Supplementary Figure 7.** The effects of NGFR deficiency on trabecular bone volume fraction, cartilage, and synovitis at 3 months after DMM. **a** and **b**, trabecular bone volume fraction of femurs in the operated legs (with OA) of female (a, n = 5) and male (b, n = 7) mice at 3 months after DMM. **c**, OARSI scoring of histology results of OA joints at 3 months after DMM. Mann–Whitney test. **d**, Synovitis score of the OA joints at 3 months after DMM. Mann–Whitney test. n = 12.

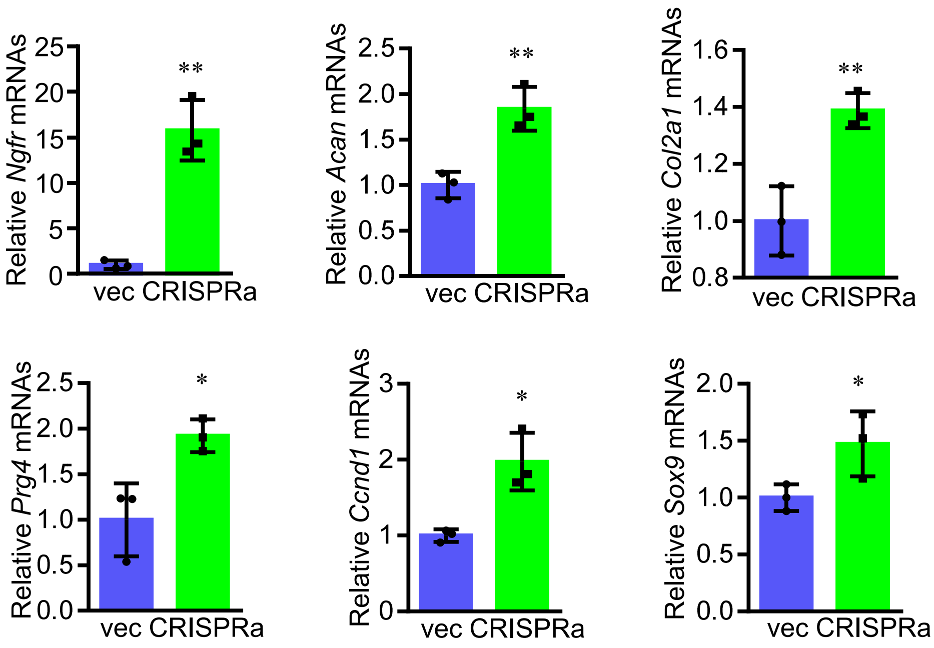


**Supplementary Figure 8.** NGFR gain-of-function upregulates osteochondral genes. Quantitative RT-PCR of mRNAs as indicated in mouse CD45$^-$ BMSCs in which NGFR activation was induced by CRISPRa. n = 3. *p<0.05, **p<0.01.

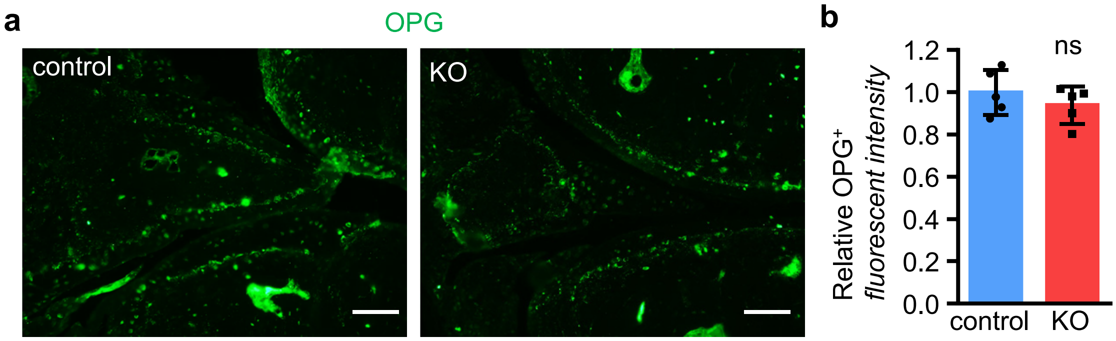


**Supplementary Figure 9.** NGFR deficiency did not upregulate OPG in skeletal cells under inflammation. **a** and **b**, Representative images and quantification of OPG IHC results. Scale bar: 80 μm. n = 5.

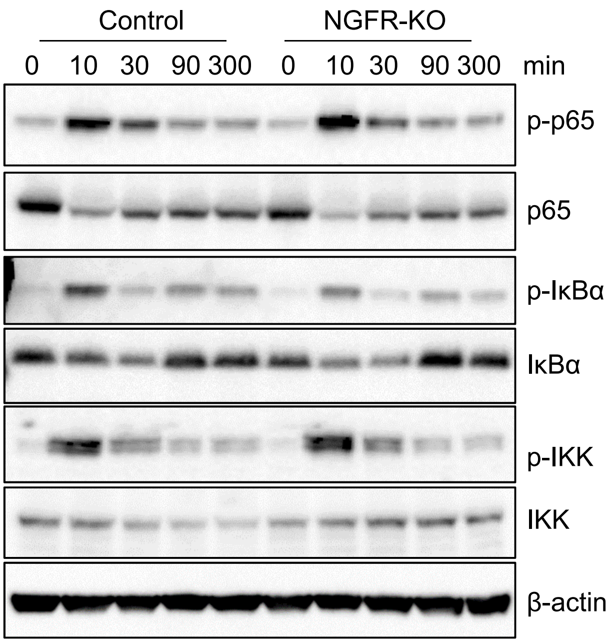


**Supplementary Figure 10**. NGFR KO hyper-activates NF-κB in skeletal cells under inflammation. The cells were treated with TNF-α for different time durations to identify the time point when maximum activation of NF-κB occurred.

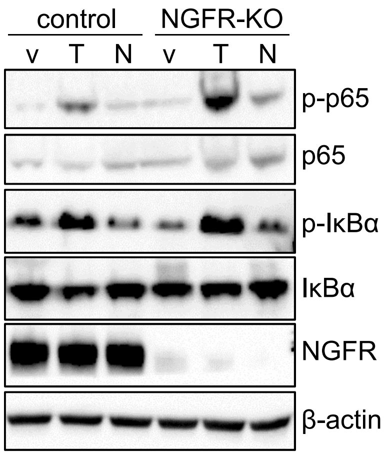


**Supplementary Figure 11.** NGFR KO hyper-phosphorylated p65 and IκBα in skeletal cells. C2C12 cells were treated with TNF and/or NGF for 15 min and then subjected to western blots. v, vehicle. T, TNF. N, NGF.

**Figure 1d-h**

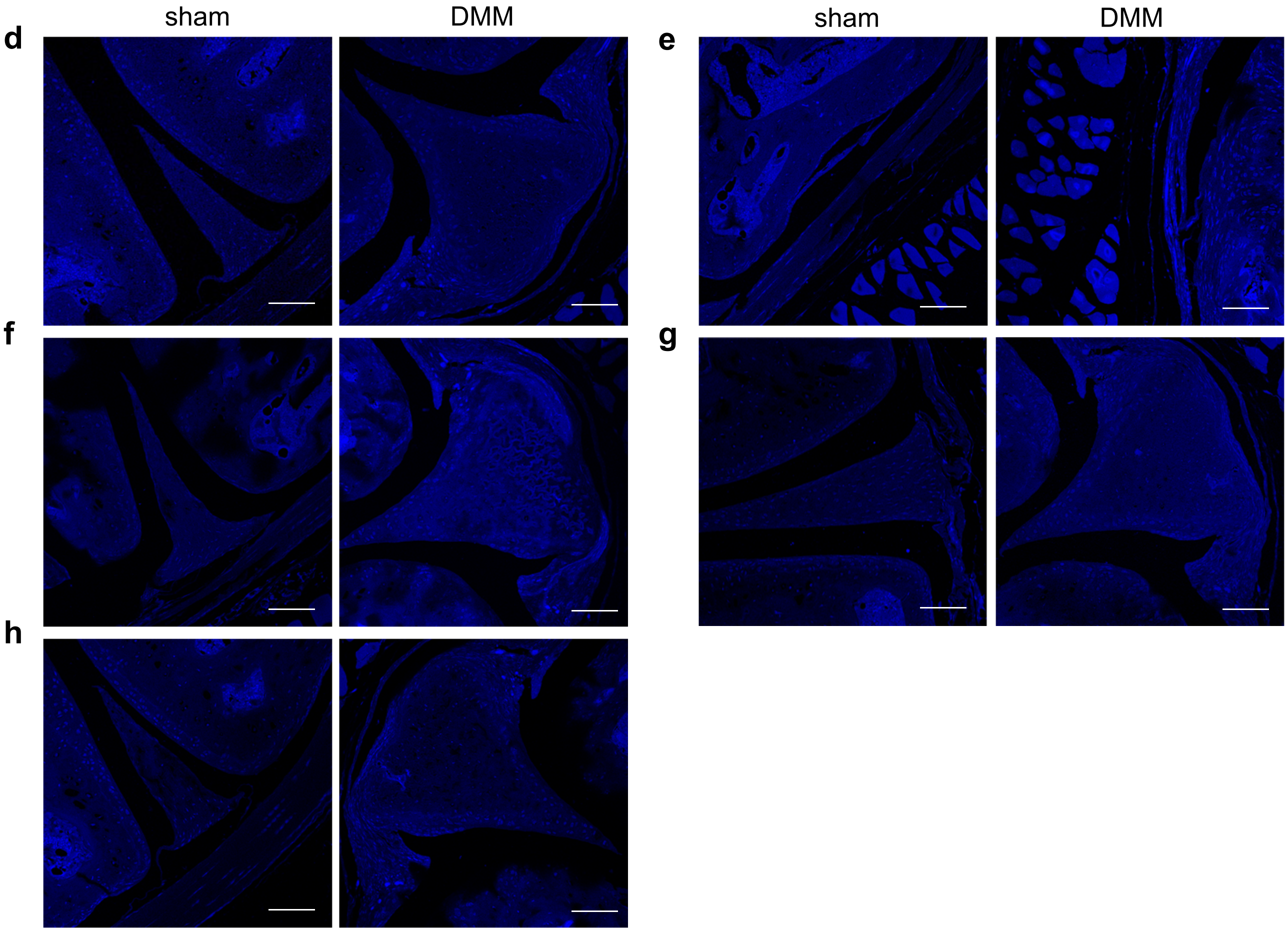


**Supplementary Figure 12.** DAPI staining for Figures 1d-h. Scale bar, 100 μm.

**Supplementary Figure 3a**

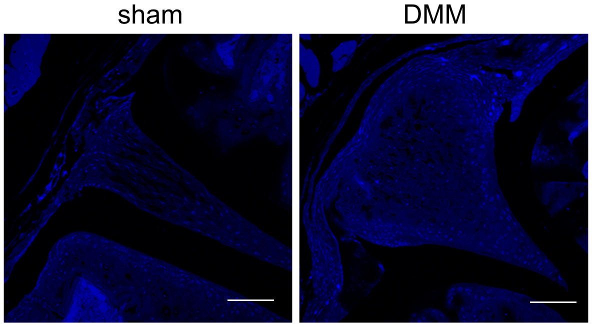


**Supplementary Figure 13.** DAPI staining for Supplementary Figure 3a. Scale bar, 100 μm.

**Figure 4b**

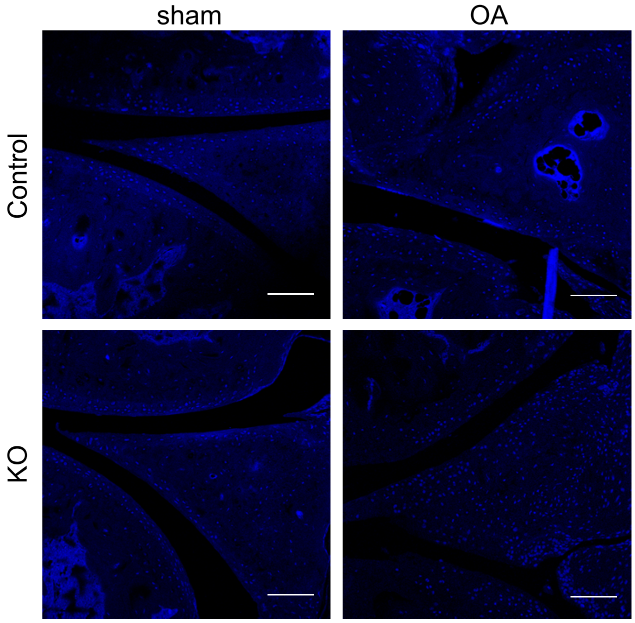


**Supplementary Figure 14.** DAPI staining for Figure 4b. Scale bar, 100 μm.

**Figure 5g**

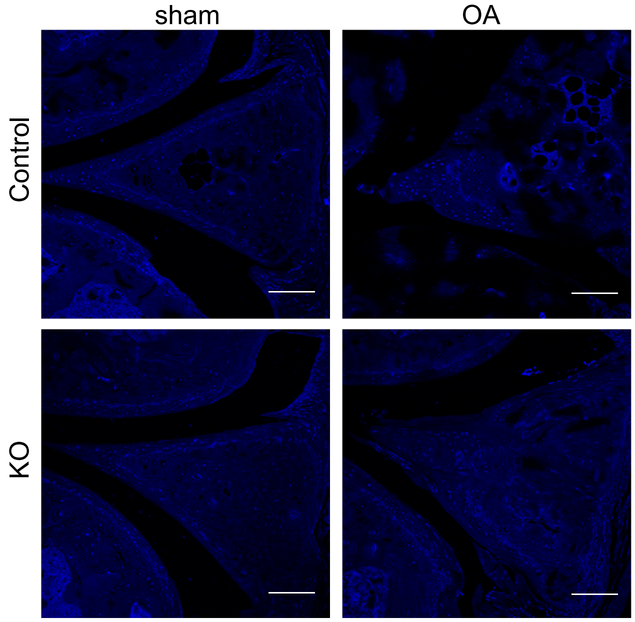


**Figure 5h**

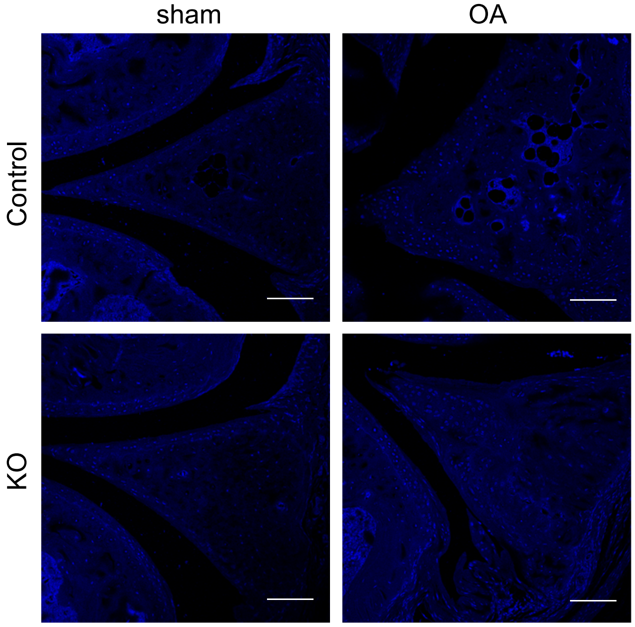


**Supplementary Figure 15.** DAPI staining for Figure 5. Scale bar, 100 µm.

**Figure 6h**

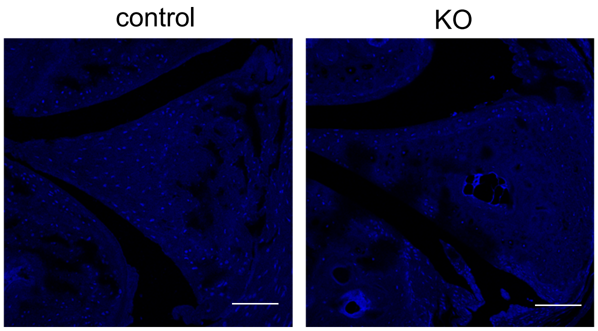


**Supplementary Figure 16.** DAPI staining for Figure 6h. Scale bar, 100 μm.
